# A mechanistic simulation of molecular cell states over time

**DOI:** 10.1101/2023.02.23.529720

**Authors:** Rossin Erbe, Genevieve Stein-O’Brien, Elana Fertig

## Abstract

Computer simulations of cell behaviors and dynamics allow for investigation of aspects of cellular biology with a ground truth that is currently difficult or impossible to generate from experimentally generated profiling data. Here, we present a mechanistic simulation of cell states that models the stochastic interactions of molecules revealing the DNA accessibility, RNA expression, and protein expression state of a simulated cell and how these states evolve over time. By designing each component to correspond to a specific biological molecule or parameter, the simulation becomes highly interpretable. From the simulated cells generated, we explore the importance of parameters such as splicing and degradation rates of genes on RNA and protein expression, demonstrating that perturbing these parameters leads to changes in long term gene and protein expression levels. We observe that the expression levels of corresponding RNA and proteins are not necessarily well correlated and identify mechanistic explanations that may help explain the similar phenomenon that has been observed in real cells. We evaluate whether the RNA data output from the simulation provides sufficient information to reconstruct the underlying regulatory relationships between genes. While predictive relationships can be inferred, direct causal regulatory relationships between genes cannot be reliably distinguished from other predictive relationships between genes arising independently from a direct regulatory mechanism. We observe the same inability to robustly distinguish causal gene regulatory relationships using simulated data from the simpler BoolODE model, suggesting this may be a limitation to the identifiability of network inference.

## Introduction

To understand the interplay between temporal changes from molecular interactions and the limitations of experimental data, mathematical simulations of molecular cell states can be useful to provide a known ground truth (Hill et al. 2016). In addition to providing benchmarks for the performance of network inference methods, using simulated genomics data generated from mathematical models can elucidate the kinds of temporal and regulatory dynamics that are likely to exist biologically. Moreover, simulations enable a broader range of evaluation of conditions and the requirements of datasets for inference as they can generate data under conditions that are infeasible or prohibitively expensive to generate experimentally. For example, simulations can readily provide temporally resolved multi-omics data from the same cells and low noise data. Thus, simulations can allow us to understand the impact of not capturing these dimensions in profiling methods on the resulting data and may illustrate cellular dynamics that are difficult to resolve from real cells.

A variety of simulation methods that attempt to generate data approximating cellular molecular states have been proposed (Pratapa et al. 2020), (Das and Mitra 2021), (Herbach et al. 2017), (Hill et al. 2016), (Gorin et al. 2022), (Thornburg et al. 2022). However, most methods do not use a direct mechanistic model with biologically interpretable parameters to simulate molecular states over time, instead parameterizing the simulation with a mathematical model such as a set of ordinary or stochastic differential equations. Additionally, most simulations do not cover all of DNA accessibility, RNA expression, and protein expression. To our knowledge, both of these features have not been implemented in the same method in previous work. Therefore, we have generated software for Multi-Omic Mechanistic Simulations (MOMS) that simulates mechanistic molecular interactions using the general model displayed in Figure 1, is fully tunable regarding molecular parameters, and simulates based on the interplay between DNA accessibility, RNA expression, and protein expression states.

**Figure 1.**
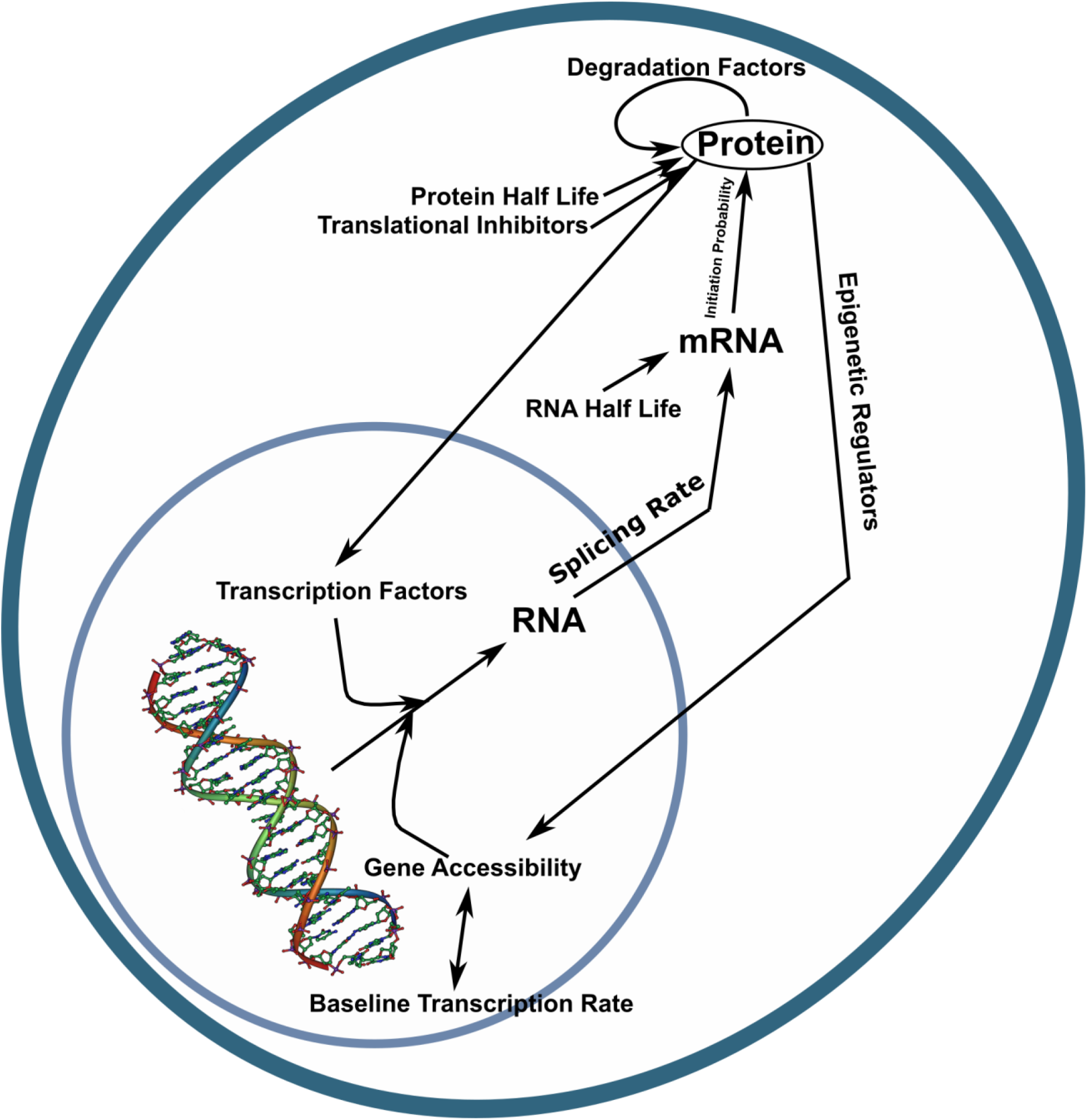
Overview of the Mechanistic Links and Parameters Incorporated in the Simulation. The simulation software proposed simulates genes with two copies that are accessible or not. The simulation accounts for baseline transcription rate and concentration of transcription factors when assessing whether transcription of a gene occurs. The RNA in the nucleus is then spliced before maturing to cytosolic mRNA which can be translated into protein, which can feedback onto the DNA as transcription factors or epigenetic regulators. Protein can additionally feedback on itself as a translational inhibitor or degradation factor. Otherwise, RNA and protein degradation occurs according to its half-life.

### A mechanistic simulation of cellular states

The MOMS simulation outputs DNA accessibility, spliced and unspliced RNA counts, and protein counts for each gene at each time point (intended to approximate a one minute time difference from the previous data output) according to the mechanistic model illustrated in Figure 1. This model uses genes as its basic units, and the cells are assumed to have diploid genomes. Any set of genes can be used to simulate cell molecular states, given a list of genes where each gene has the required parameters specified: Baseline Transcription Probability, Transcription Factors, Transcriptional Repressors, Epigenetic Upregulators, Epigenetic Downregulators, Splicing Rate, RNA half-life, Protein half-life, Translation Initiation Probability, Translational Inhibitors, and Protein Degradation Factors (Figure 1). This set of genes can be generated randomly or based on some known biological network input by the user.

Each copy of a gene can be either accessible or not accessible. For the simulation, this means the gene is either transcribed at a normal rate based on transcription factor binding (if accessible) or is much less likely to be transcribed because transcription factors cannot usually bind to the DNA (if not accessible). This is specified in the simulation in a binary manner, where each gene copy is accessible or not and there is a small (tunable) probability of transcription even when not accessible. If a gene is accessible, there is a baseline probability of the standard transcription factors binding per unit time of the simulation, which is intended to represent approximately one minute of real time. Each gene can be assigned a set of specific transcriptional activators and repressors that promote or inhibit RNA transcription, as well as epigenetic activators and repressors that can change the accessibility state. The impact of these regulatory genes is based on the concentration of the corresponding proteins. Thus, the volume of the nucleus and cytosol are specified in the simulation and the binding probability is calculated according to the equation 1 - e^-[protein]. The binding affinity is assumed to be equal for all proteins. Whether binding occurs at each unit time is evaluated with a random number generator using the specified binding probability. The splicing rate is specified for each gene and the spliced and unspliced RNA transcript counts are tracked separately. RNA degradation is controlled by the RNA half-life parameter, which is specified for each gene.

Genes in the simulation are assumed to produce mRNAs. RNA is translated to protein based on a specified probability for how likely a spliced transcript is to bind a ribosome per unit time, as well as the concentration of translational inhibitors. Translational inhibitors are protein products of specified genes. Protein degradation is determined by the protein half-life parameter specified for each gene, as well as the concentration of any protein specific degradation factors (i.e. other proteins that bind it and cause it to be degraded).

This simulation structure is probabilistic and thus the results are to some extent stochastic. As in our previous work, we select a probabilistic model as the basis of cell behavior because the precise moment molecules collide and bind is not usually predictable in advance without measuring cell state to a degree that is far beyond current technologies (Fertig et al. 2011).

Often we may want to randomly generate a gene set and its parameters rather than specifying each parameter manually. Thus, we provide functionality to generate the parameters for random gene sets, with parameter values that fall within normal biological ranges based on reference to experimental data. All parameter ranges are user tunable. For the simulations presented below, the baseline transcriptional probability was set between 0.01 and 0.001, the maximum number of transcriptional regulators was set to 3 proteins, and the maximum number of epigenetic regulators was set to 4 proteins. The splicing rate was randomly set for each gene between 5 and 10 time points (approximately 5 to 10 minutes). The RNA half-life range was between 60 to 900 time points and the protein half-life range was between 720 and 3600 time points, intended to reflect the fact that most proteins are degraded much more slowly than RNA, usually via proteasomal and lysosomal pathways (Schwanhäusser et al. 2011), (Cambridge et al. 2011), (Sha, Zhao, and Goldberg 2018). Each gene could have at most one translational inhibitor and at most two protein degradation factors. The probability of translation initiation for an unbound mRNA at each time point was set between 0.1 and 0.75.

The Julia software to run MOMS simulations is made freely available on GitHub at https://github.com/FertigLab/MOMSCellSimulations.

### Impact of parameters on molecular cell states

The outcome of a simulation is determined by the parameters provided for each gene, the initial conditions of the simulated cell, plus a random number generator that determines the outcome of stochastic elements. A key to the variation in the simulations is gaining some insight into how much impact changing the random number generator seed and the initial conditions changes the observed cell states over time.

To do this, we randomly generated the parameters for ten genes and simulated a cell for 1000 time points (Figure 2A). The simulation was then repeated, changing only the random seed used for the random number generator used to evaluate whether protein binding occurs during a particular time interval, to determine how deterministic the expression levels of RNA and protein are in the simulation (Figure 2B). We observe general similarity, but with notable differences. The lowest expressed gene (at both the RNA and protein level) has a small increase in RNA expression at the RNA level, which leads to a larger increase at the protein level for most of the simulation period. For the other genes, little to no change in expression is observed at both the RNA and protein level. This result demonstrates that while the simulation is stochastic, the distribution of values of gene expression are predictable.

**Figure 2.**
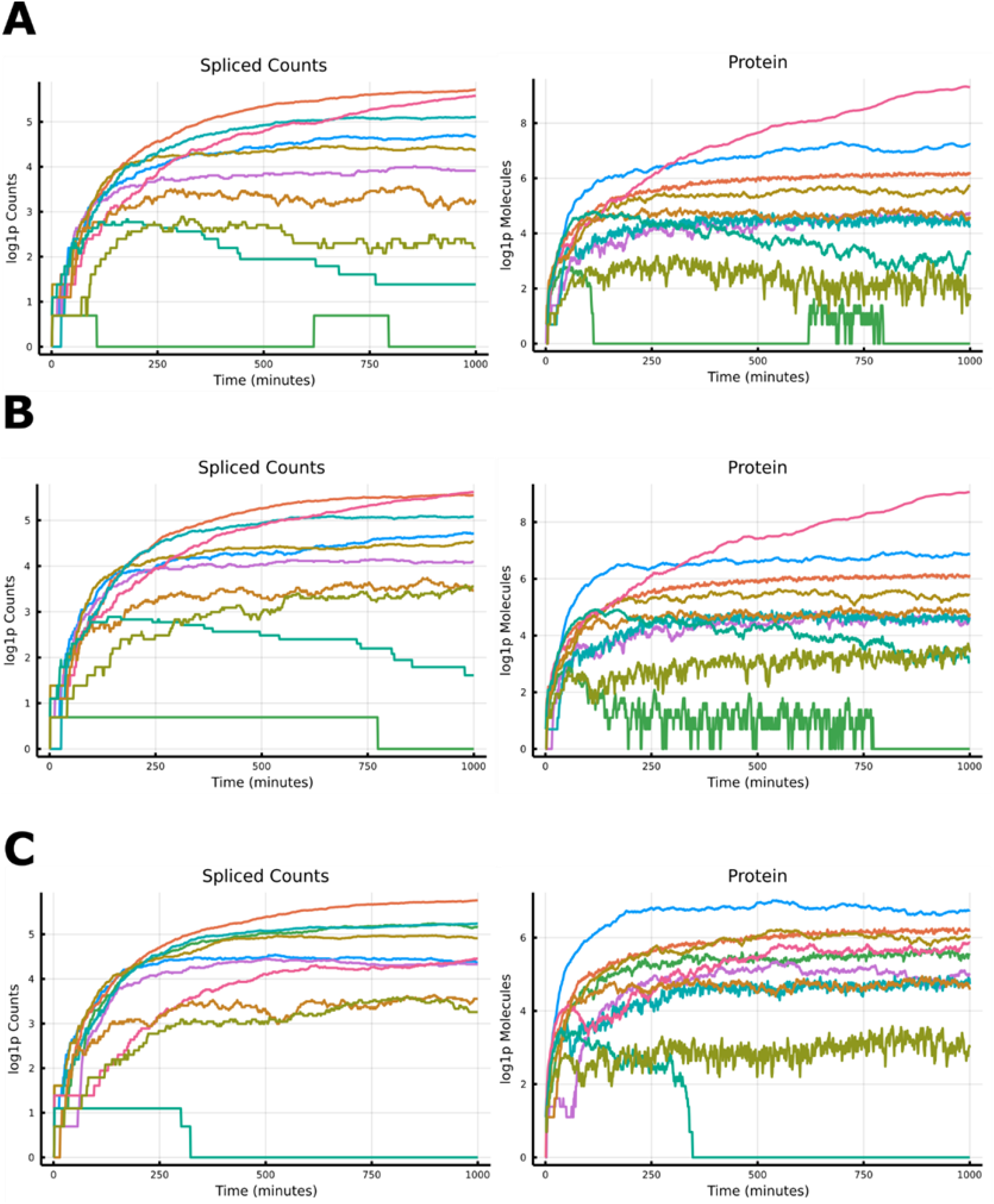
Impact of Random Seed and Initial Conditions on Simulation of RNA and Protein Levels. **A** Spliced RNA counts and protein expression of a baseline ten gene simulation over 1000 time points. **B** Spliced RNA counts and protein expression with a different random seed than A. **C** Spliced RNA counts and protein expression with a different set of initial conditions from A.

Each simulation is provided with a set of initial conditions to parameterize the interactions between genes. Each gene’s expression at the RNA and protein level begins at a random value between 0 and 2 molecules. In order to test how much impact the initial conditions of the simulation had on the molecular state in the long run, we generated a different set of initial conditions randomly for the same gene set used in Figure 2A. We then ran the simulation with the new initial conditions for 1000 time points (Figure 2C). Here, the expression of several genes changes substantially. Notably, the pink labeled protein, which is highest expressed in Figure 2A is three orders of magnitude lower expressed with different initial conditions. The lowest expressed green labeled gene from 3.2A is now expressed much more highly, at about the same level as the pink labeled gene. The turquoise labeled gene’s expression also goes to zero at the RNA and protein level by the halfway mark in the simulation and does not make a resurgence. The other genes’ expression remains similar to the previous initial conditions. These results indicate that the initial conditions can have a large impact on future cell states, even if the levels of all genes’ expression are low initially.

With a mechanistic model of cell states that contains parameters corresponding to molecular mechanisms, we can perturb those parameters to attempt to understand the impact those parameters have on cellular states. Using the same baseline set of genes used in Figure 2A, we perturbed the splicing rate of each gene and reran the simulation (Figure 3A). Changing the splicing rate had some impact on the early states of the simulation, particularly at the RNA level, but in the long run very little difference in expression at the RNA or protein level is observed with the different splicing rates, which may indicate that splicing rate does not have a large impact on RNA and protein expression levels long term, at least when the splicing rate is only a small fraction of the half-life of the molecules (as is usually the case in real cells) (Alpert, Herzel, and Neugebauer 2017).

**Figure 3.**
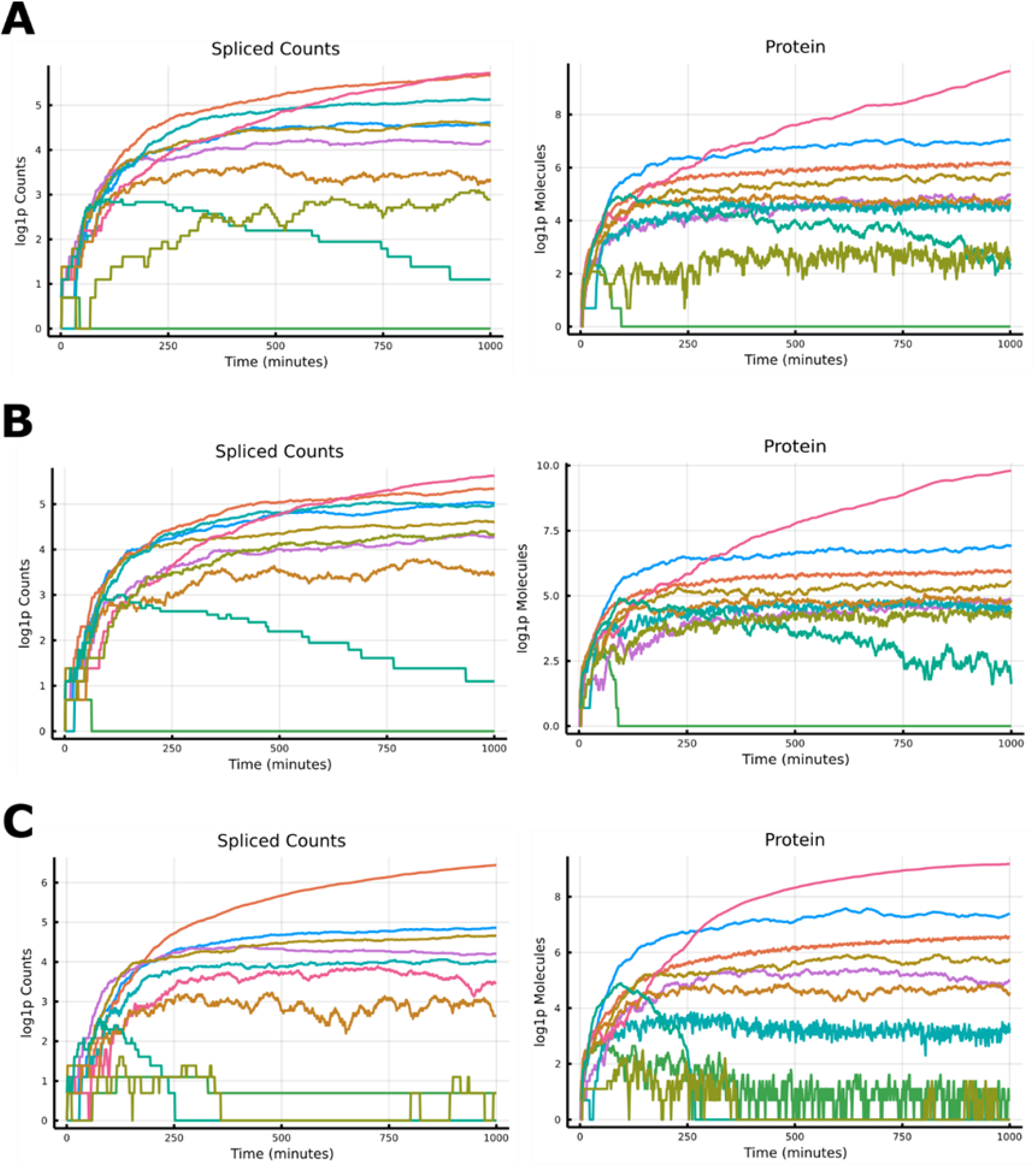
Impact of Changing Splicing Rate and half-life Parameters. **A** Spliced RNA counts and protein expression after changing splicing rate of each gene. **B** Spliced RNA counts and protein expression after changing protein degradation rate of each gene. **C** Spliced RNA counts and protein expression after changing RNA degradation rate of each gene.

Another important parameter is RNA and protein half-life. We observe that perturbing protein half-life mostly yields a large change in protein expression of one gene, the pink gene, which is even higher expressed (Figure 3B). Only minor changes to RNA expression are observed, as might be expected. Perturbing RNA half-life leads to three genes having very low RNA expression levels. Protein expression of these genes swiftly follows suit, leaving those three genes at or near zero expression for the remainder of the simulation (Figure 3C).

### Effects of gene perturbations on cellular states

Limitations to temporal profiling technologies currently make it difficult to track the impact of perturbations within the same single cell over time. We can evaluate how perturbations may impact cell states over time using simulated data. To evaluate the way a gene knockout impacts future expression states, we modified the simulation from Figure 2A. We allow the simulation to proceed for fifty time points and then allow no more of the RNA transcripts of the blue gene to be produced, mimicking the result of a fully deleterious mutation that leads to immediate RNA degradation (Figure 4A). The remaining RNA degrades rapidly, while the remaining protein degrades over the next ~150 time points. This knockout leads to substantial changes in gene expression, particularly at the protein level. The protein expression of the pink gene reaches its maximum three orders of magnitude lower than in the original simulation, while the tan gene ends three orders of magnitude higher. Two gene’s protein expression falls to zero, three stay relatively constant, and two others increase in expression about tenfold. Interestingly, only two of the observed changes in protein expression appear to be driven by changes in RNA expression level: the knockout gene and the orange gene. The other differences observed appear to be mediated via the direct regulation of proteins by proteins (both through degradation by inhibiting translation).

**Figure 4.**
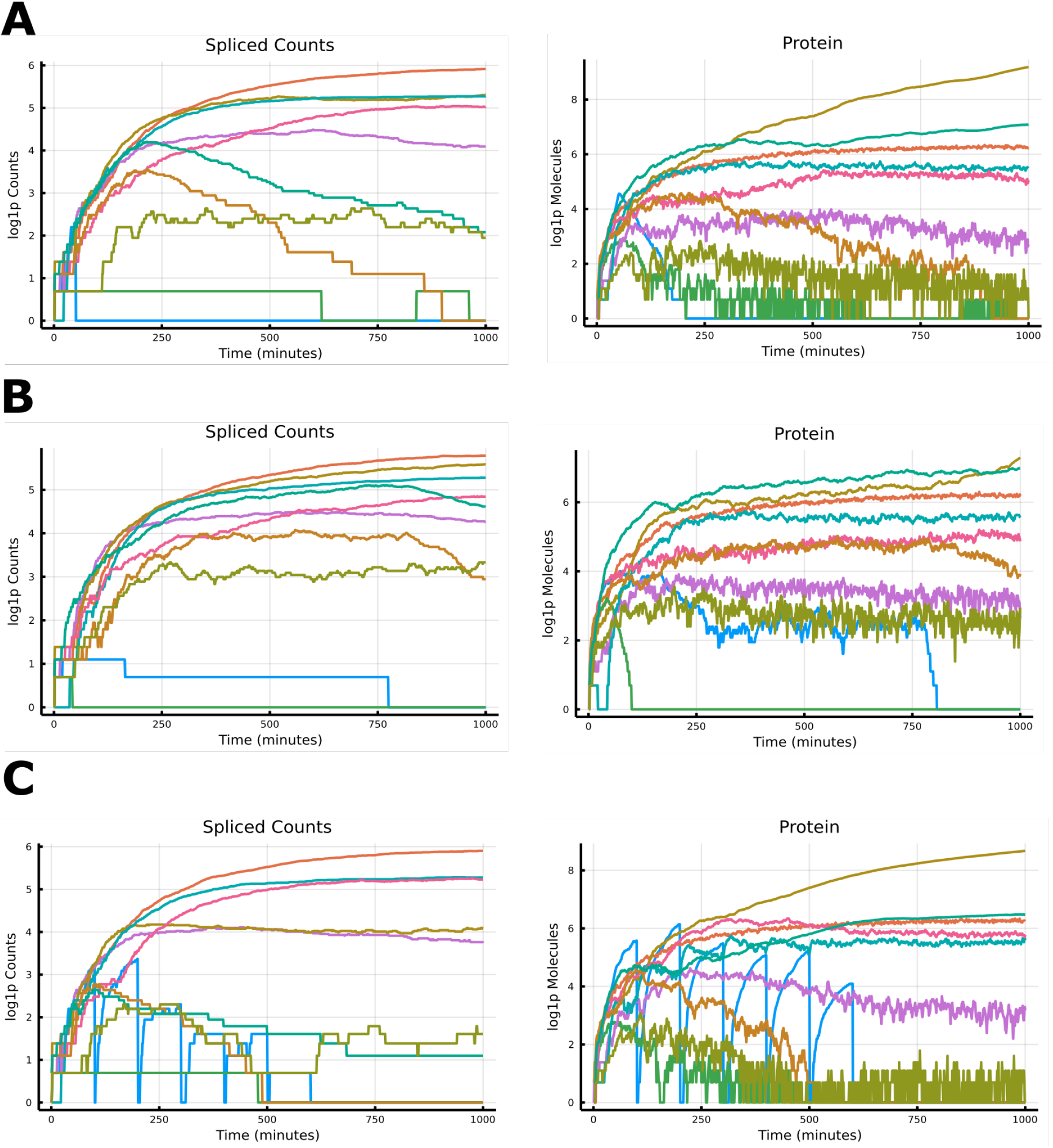
Impact of Perturbations of RNA and Protein Expression Levels over Time. **A** Spliced RNA counts and protein expression if gene 1 (blue) is set to an epigenetically repressed state after t=50. **B** Spliced RNA counts and protein expression if gene 1 (blue) is prevented from transcribing after t=50. **C** Spliced RNA counts and protein expression if gene 1 (blue) is set to zero every 100 time points.

We compared this knockout simulation to perturbing the same gene at an epigenetic level. After time t=50, the blue gene was set to a non-accessible state (Figure 4B). The RNA expression slowly decreases over time as the remaining counts degrade, while a very low level of expression is allowed to occur despite general gene inaccessibility. Protein expression slowly falls as well, not reaching zero until 800 time points in. The slower change means the change in expression levels plays out slightly differently. The pink gene’s protein expression falls to a similar level as in the knockout, however, the tan gene does not reach as high of an expression level and has similar expression as the turquoise gene, which is again increased in expression from the original simulation to the same level. The RNA expression of the orange gene does not start to fall until the last two hundred time points, leaving both its RNA and protein expression at close to the original level by the end of the simulation.

We can also investigate a wider variety of perturbations than can be performed in real cells using current technologies. For example, we can set the RNA expression level of a gene to zero instantaneously and then trace how each cell responds over time at an epigenetic, transcriptional, and proteomic level. In order to test what this sort of perturbation would look like, we created a simulation using the same gene set as Figure 2A and at time t = 100 we set the spliced RNA counts to zero. We repeated this perturbation every 100 time points through the end of the simulation. We observe that the expression of the gene rapidly returns to its original value after the first perturbation. This result suggests that the molecular state of the cell is able to reestablish the RNA expression of an upregulated gene, even in the absence of any RNA from that gene (Figure 4C). Despite rebounding in expression level five times, the expression profiles of the other genes in the simulation quickly reach approximately the same levels they occupy in the knockout simulation, which may indicate that even temporary loss of the blue gene’s expression immediately begins moving the cells towards the state observed in the knockout simulation, while temporary reexpression of the blue gene does not restore the previous state.

### Correlation of RNA expression and corresponding protein expression varies substantially

The naïve expectation is that an mRNA and the corresponding protein will in most cases have strongly correlated expression values. However, due to the time lag between transcription and translation as well as due to the influence of other regulatory factors, this expectation will not necessarily hold. Profiling studies suggest that in many cases, an mRNA and its corresponding protein are almost entirely uncorrelated (Gry et al. 2009), (de Sousa Abreu et al. 2009), (Vogel and Marcotte 2012). In order to investigate this phenomenon, 100 simulations with different gene sets were generated and the correlation between the RNA and protein levels was assessed. The mean correlation between RNA and protein expression was 0.63, with a median of 0.80 (Figure 5A). The correlation coefficient for a substantial minority (10.8%) of genes was less than zero. Often, we see this occur because of the influence of another gene. For example, if gene 1 is strongly regulated at the protein level by gene 2 then gene 1 RNA and gene 1 protein may not be correlated because the primary determinant of gene 1 protein levels may be gene 2 protein levels. We also observe that some regulatory networks result in genes and proteins with higher correlations than others. Median correlation varies substantially between simulations randomly assigned different gene-level parameters as described previously (Figure 5B), with some simulations above 0.9 and others below zero. The simulations with median correlation below zero often have one or two genes that strongly regulate the protein levels of many other genes. These one or two genes are usually very highly expressed and thus largely make the expression level of other genes irrelevant to cell state due to the overwhelming influence of one to two genes. It is important to note that this low correlation is observed even with “perfect measurement” of the RNA and protein levels; without the measurement noise that would exist with any profiling technique used to measure the expression levels in real cells.

**Figure 5.**
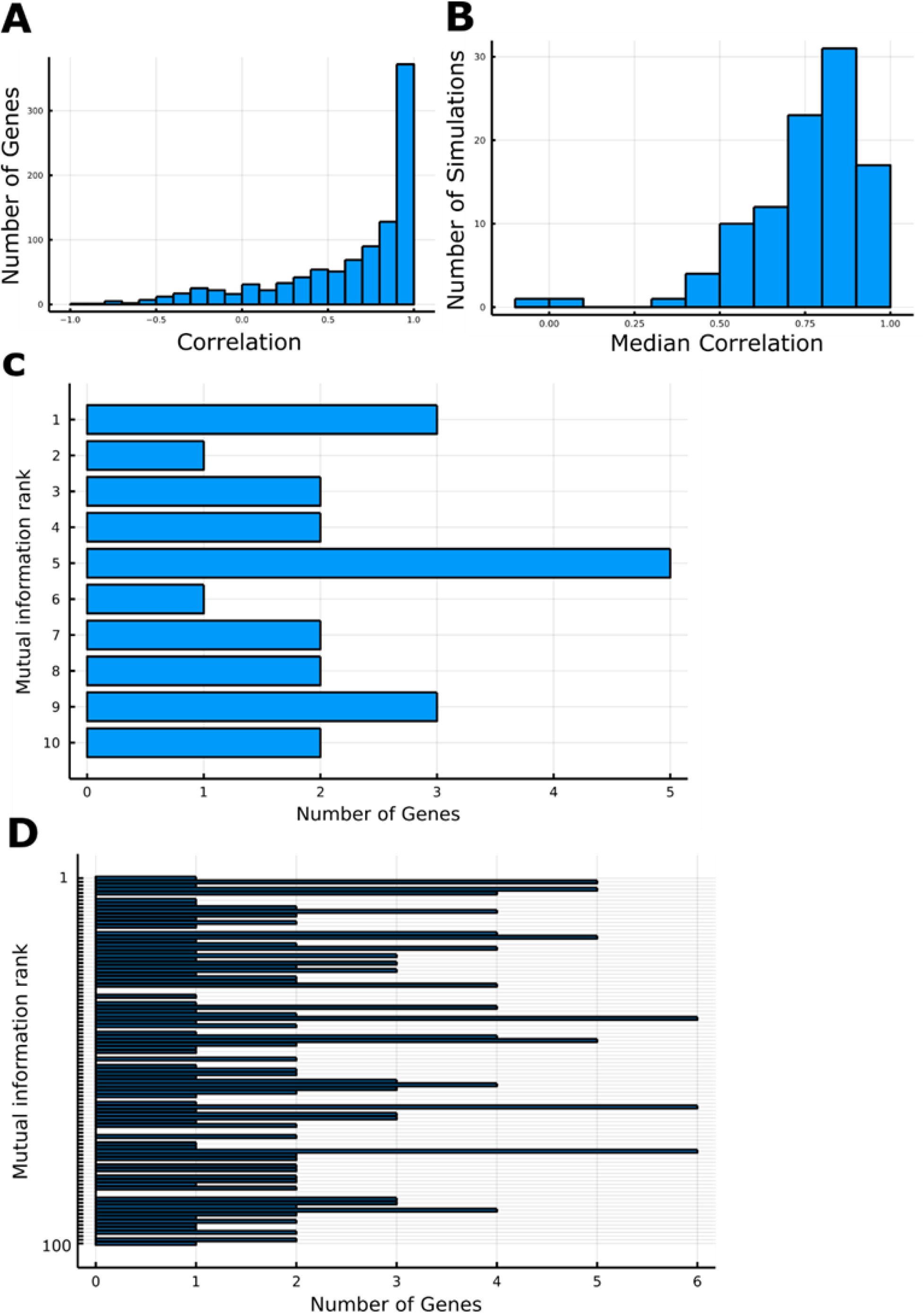
Correlation and Information between RNA and Proteins. **A** Histogram of the correlation of each corresponding RNA-protein pair across 100 ten gene simulations. **B** Histogram of the median correlation of each simulation between each corresponding RNA-protein pair across 100 ten gene simulations. **C** Histogram of the mutual information rank among all genes provided by each causal gene pair among 100 ten gene simulations. **D** Histogram of the mutual information rank among all genes provided by each causal gene pair among 100 hundred gene simulations.

### The mutual information between the RNA expression of two genes is insufficient to reliably distinguish causal gene pairs

The goal of many genomics analyses is to extract mechanistic understanding from the data (Erbe et al. 2022). To accomplish this, many gene network inference methods have been developed, which attempt to infer causal regulatory relationships between genes using gene expression data (Margolin et al. 2006), (Chan, Stumpf, and Babtie 2017), (Huynh-Thu et al. 2010), (Osorio et al. 2020), (Matsumoto et al. 2017), (Papili Gao et al. 2018), (Deshpande et al. 2019), (Qiu et al. 2020). Despite the myriad of approaches, independent assessments indicate that these methods often cannot robustly predict known causal interactions between genes (Chen and Mar 2018), (Pratapa et al. 2020), (Stone et al. 2021). In order to better understand the challenge facing these network inference methods, we simulated 2000 cells with the same regulatory parameters and randomly selected an expression profile for each cell over 500 simulated time points. The resulting matrix mimics the output of single cell RNA-seq data, without any measurement noise. We performed this simulation experiment with both ten gene cells and 100 gene cells. We then used the resulting count matrix to find the mutual information each gene pair provided and selected the gene pairs with a causal regulatory interaction between them. The mutual information for causally interacting gene pairs was not significantly higher than non-causal genes for the ten or 100 gene data set (Figure 5C-D). This result indicates that the statistical relationships between genes that are not directly causal prevent differentiation of causal and non-causally related genes on the basis of predictive power, even in a noiseless simulation. While the rank distribution in the ten gene set is slightly shifted towards lower ranks, this shift is not nearly sufficient to reliably distinguish these gene pairs from non-causal ones.

In order to determine whether this result was based on the specifics of our simulation model or would also occur in other biologically-driven simulation frameworks models, we assessed whether the information between causally related genes was higher when using the BoolODE simulation (Pratapa et al. 2020). BoolODE takes a causal regulatory network as input and uses the regulatory relationships to parameterize a set of ordinary or stochastic differential equations to simulate single-cell gene expression values. We simulated 2000 cells from two different causal regulatory networks provided by BoolODE (Pratapa et al. 2020): one derived from studies of Gonadal Sex Determination (GSD) and the other designed to produce a trifurcation in cell lineage. In both of these simulations, the mutual information between a pair of genes was again insufficient to reliably distinguish causal gene pairs (Figure 6). While the GSD simulation does show a slightly higher proportion of causal gene pairs with high mutual information ranks, many of the causal gene pairs are still very low ranked, preventing robust causal inference for these gene pairs. The trifurcation simulation shows very little bias in rank distribution for the causal genes pairs, indicating there is nearly no signal distinguishing these gene pairs from random gene pairs. Taken together, the results from the MOMS simulation and BoolODE simulation suggest direct causal inference from single-cell RNA-seq datasets may not be possible for a large subset of genes because the statistical relationship between expression values does not distinguish causally related and correlated gene pairs even in the clean simulated cases presented here.

**Figure 6.**
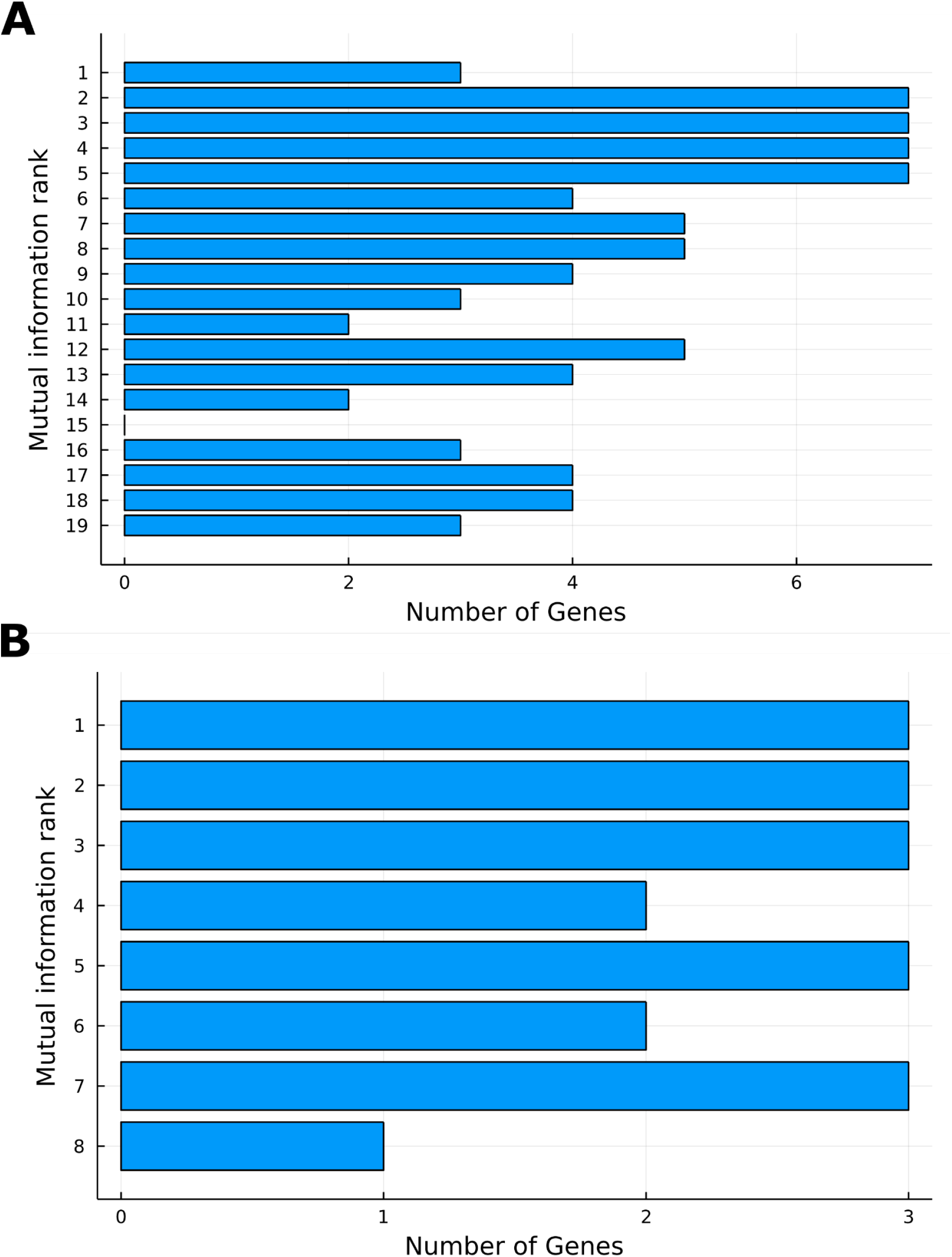
Mutual Information Between Causal Gene Pairs in BoolODE. **A** Histogram of the mutual information rank of causally related gene pairs compared to all gene possible gene pairs for each gene in the BoolODE GSD simulation across 2000 cells. **B** Histogram of the mutual information rank of causally related gene pairs compared to all gene possible gene pairs for each gene in the BoolODE trifurcation simulation across 2000 cells.

### Gene accessibility provides limited predictive information about RNA and protein expression levels except in a fully inaccessible state long term

The chromatin accessibility of a gene and its regulatory regions controls the capacity for transcription factor binding, and subsequent gene transcription (Klemm, Shipony, and Greenleaf 2019). While there is evidence that chromatin accessibility is better described as a continuum than a dichotomy (Poirier et al. 2008), the current state of the art in large-scale profiling with single-cell ATAC-seq methods only provides a binary readout, with the DNA fragments in a cell either being sequenced or not (Buenrostro et al. 2015). For the sake of simplicity, and to conform to the type of data generated by single-cell ATAC-seq, we use a binary version of chromatin accessibility in the MOMS simulation.

While the impact of long term repressive impact of chromatin inaccessibility is generally agreed upon, and is recapitulated by the MOMS simulation (Figure 4B), the impact of variation in chromatin accessibility on gene expression is not well established. Using the MOMS simulation, we explored the mutual information between gene accessibility and gene expression across 100 different ten-gene and hundred-gene simulations to determine the impact of variation in chromatin accessibility on the expression of the regulated gene. Across both the ten and hundred gene simulations, the average mutual information between gene accessibility and the expression level of the corresponding gene at RNA or protein level was much lower than the mutual information between RNA and protein from the same gene (Figure 7). The main reason for this discrepancy in information content appears to be the binary nature of the accessibility states, allowing only a representation corresponding to neither gene copy accessibility, one gene copy accessible, or both gene copies accessible. Thus, the much more variable RNA-protein dynamics can contain much more information. Additionally, we find that in the simulated representation of a single-cell data set introduced in the previous section, variance in gene accessibility state is almost entirely uncorrelated with variance in spliced RNA expression (r = 0.057). However, we observe a somewhat more moderate correlation with variance in protein expression (r = 0.40)

**Figure 7.**
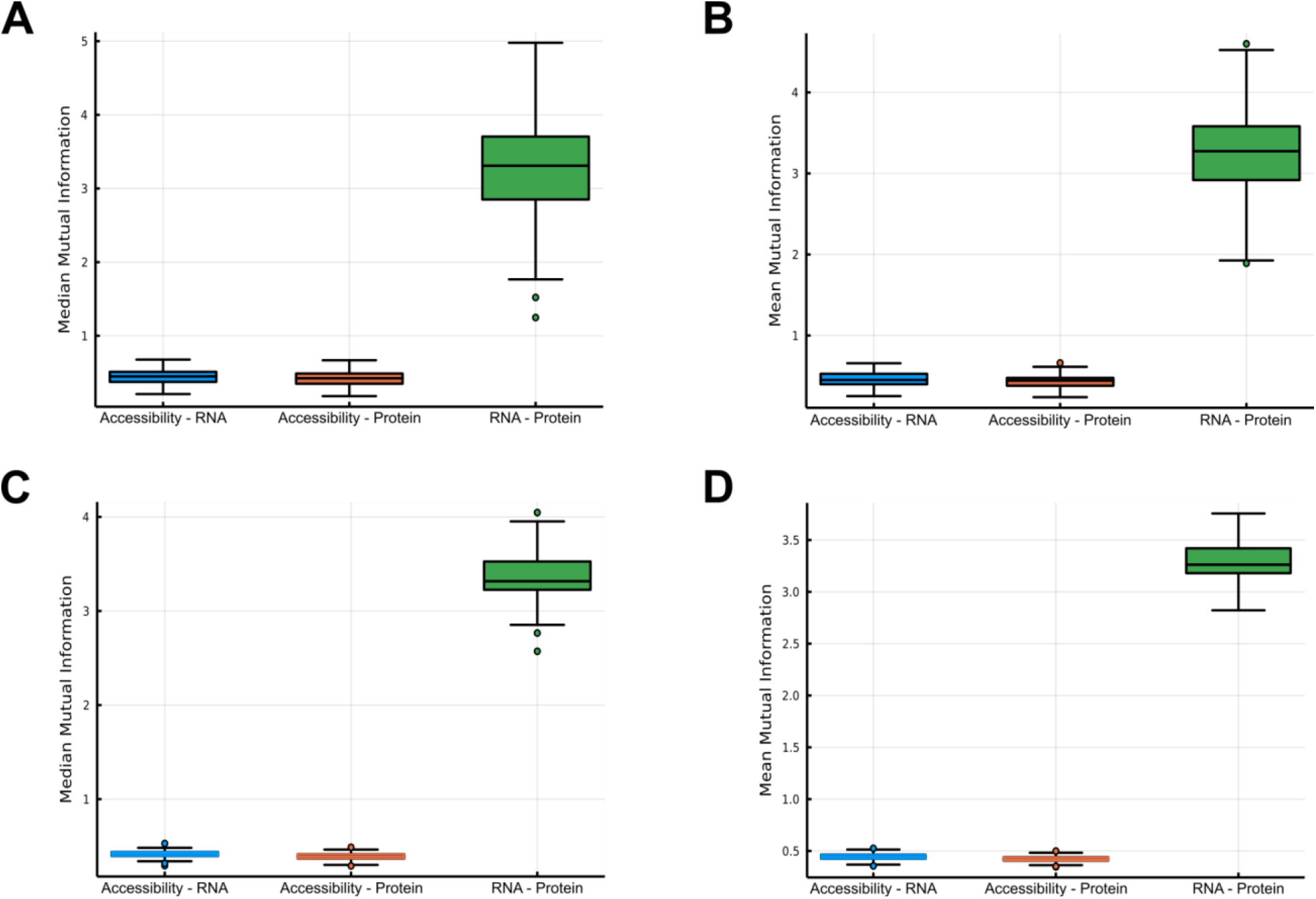
Mutual Information Between Simulated Genomics Types. **A** Boxplots of the median mutual information from 100 ten-gene simulations comparing the mutual information between accessibility of a gene and its RNA expression, accessibility and protein expression, and RNA expression and protein expression. **B** Boxplots of the mean mutual information from 100 ten-gene simulations comparing the mutual information between accessibility of a gene and its RNA expression, accessibility and protein expression, and RNA expression and protein expression. **C** Boxplots of the median mutual information from 100 hundred-gene simulations comparing the mutual information between accessibility of a gene and its RNA expression, accessibility and protein expression, and RNA expression and protein expression. **D** Boxplots of the mean mutual information from 100 hundred-gene simulations comparing the mutual information between accessibility of a gene and its RNA expression, accessibility and protein expression, and RNA expression and protein expression.

## Conclusion

We present a mechanistic simulation of cell states across epigenetic, transcriptomic, and proteomic cell states, MOMS. MOMS includes parameters for splicing rate and degradation of genes, allowing the impact of these to be assessed. The simulations we have presented indicate that degradation rate parameters are substantially more impactful on the long term RNA and protein expression rate than splicing rate, though perturbations of either parameter have observable impacts on expression.

We examined the relationships between the expression values of RNAs and their corresponding proteins in order to determine if the proposed simulation matched reports from cells of frequent low correlations between these pairs. We did find a wide array of correlations between corresponding RNA and protein pairs, including many near or even below zero. This result suggests that our simulation captures important regulatory features that give rise to those dynamics *in vivo*. Additionally, the simulation provides explanations for how this phenomena can occur. Frequently, we find it is observed due to the overwhelming influence of another gene that obfuscates the relationship because protein or RNA expression is most strongly influenced by the other protein.

One limitation of MOMS is that it supplies output at discrete time intervals rather than allowing for continuous assessment of cell states. However, this choice allows for a highly computationally efficient simulation relative to a continuous model. For the purpose of allowing many simulations across many cells to be performed under many different perturbation conditions, we believe this tradeoff will often be worthwhile.

We further investigated the ability to distinguish correlation from causation using simulated RNA data. By randomly sampling cells from simulations over time (to approximate how scRNA-seq captures single time points of RNA expression) we find that causal gene pairs often have lower mutual information between them than other non-causal gene pairs. We further validated this result using a different type of simulation, BoolODE (Pratapa et al. 2020). Taken together, the inability to distinguish causal from non-causal gene pairs even in simulated data sets with a ground truth suggests robust causal gene network inference using only single cell RNA-seq count matrices as input may be subject to pervasive inaccuracies.

## Notes

https://github.com/FertigLab/MOMSCellSimulations

### Competing Interest Statement

The authors have declared no competing interest.

